# Mechanically activated bone cell derived extracellular vesicles regulate angiogenesis in a manner that is dependent on the stage of lineage commitment

**DOI:** 10.1101/2025.07.29.667365

**Authors:** C.S. Martins, M. Maggio, C. Gorgun, M.Y. Brunet, M. Dobricic, R. Almasri, F. J. O’Brien, L. O’Driscoll, D. A. Hoey

**Affiliations:** Trinity Centre for Biomedical Engineering, Trinity Biomedical Sciences Institute, Trinity College, Dublin, D02 R590, Ireland; Dept. of Mechanical, Manufacturing, and Biomedical Engineering, School of Engineering, Trinity College Dublin, Dublin D02 DK07, Ireland; School of Pharmacy and Biomolecular Sciences, Royal College of Surgeons in Ireland, Dublin, D02 VN51, Ireland; Tissue Engineering Research Group, Department of Anatomy and Regenerative Medicine, Royal College of Surgeons in Ireland, Dublin, D02 YN77, Ireland; School of Pharmacy and Pharmaceutical Sciences, Trinity Biomedical Sciences Institute, & Trinity St. James’s Cancer Institute, Trinity College Dublin, Dublin, D02 R590, Ireland; Advanced Materials and Bioengineering Research Centre, Trinity College Dublin & Royal College of Surgeons in Ireland

## Abstract

Bone regeneration requires a finely tuned interplay between osteogenesis and angiogenesis. While current treatments such as auto/allografts, provide support, they often fail to promote adequate vascularisation necessary for complete repair. Extracellular vesicles (EVs), as mediators of intercellular communication, have emerged as promising acellular nanotechnologies for tissue regeneration due to their bioactive cargo and low immunogenicity. Mechanical stimulation, a known enhancer of bone cell function, can modulate EV cargo and potentially improve regenerative efficacy. In this study, we investigated how mechanical stimulation, and the stage of mesenchymal lineage commitment influence the angiogenic potential of secretomes and EVs derived from mesenchymal stromal/stem cells, osteoblasts, and osteocytes. Our findings reveal that both cell mechanical stimulation and their differentiation stage significantly modulate the angiogenic properties of the resulting EVs. Among the tested conditions, mechanically-stimulated osteocyte-derived EVs demonstrate superior angiogenesis, promoting endothelial cell migration, tube formation, and CD31 expression. These effects were further validated in a pre-clinical *ex ovo* chick chorioallantoic membrane assay, where robust neovascularisation was observed. This work highlights the critical role of both mechanical cues and cell differentiation stage in regulating the angiogenic capacity of EVs and proposes mechanically activated osteocyte-derived EVs as a novel pro-angiogenic nanotherapeutic for bone repair.

## Introduction

Bone regeneration remains a significant clinical challenge, as many defects such as complex fractures, tumour resections, and tissue necrosis cannot heal without clinical intervention [1]. Current treatments, including the gold standard of auto/allografts [2], have several limitations such as the constrained donor sites and supply, the possibility of an immune rejection, and the need for additional surgery [3]. Furthermore, most current therapies fail to fully support the vascularisation necessary for optimal bone regeneration [4, 5]. Bone is a highly dynamic tissue that relies on a well-developed vascular network for its formation, maintenance, and repair [6–8]. Blood vessels not only supply oxygen and nutrients but also serve as a signalling hub that regulates bone morphogenesis and homeostasis. Specifically in bone repair, the formation of new blood vessels allows for the migration of immune cells into the defect site, as well as osteoprogenitor cells. An important aspect of both bone morphogenesis and repair is the coupling between angiogenesis, the formation of new blood vessels, and osteogenesis, the formation of new bone [9, 10]. Within bone vasculature, type H vessels play a critical role in this coupling and are characterised by high expression of CD31 and Endomucin (CD31+, EMCN+) [11]. These vessels form a niche for RUNX2+ osteoprogenitors, where both endothelial cells and osteoprogenitor cells are communicating with each other, through secreted factors such as vascular endothelial growth factor (VEGF) and bone morphogenetic proteins (BMPs) [12]. This coupling and cross communication is essential to maintain bone homeostasis and facilitate effective repair[10, 13].

A key biological mechanism of cell-cell communication is via extracellular vesicles (EVs). EVs are lipid bound nanoparticles released by cells that contain diverse cargoes such as DNA, RNA, and proteins, many of which have been shown to be modulate both angiogenesis and osteogenesis [14–17]. They have also demonstrated potential as an alternative to cell-based therapies, as they have increased stability and low immunogenicity [18, 19]. The cargo of the EVs can be tuned by conditioning the parent cell in order to enhance their regenerative capacity. Mechanical stimulation is a known positive regulator of bone physiology and repair *in situ* and it has been shown *in vitro* that mechanical stimulation of the parent cell can modulate cargo within the secreted EVs [20–24]. Specifically, our lab has previously shown the impact of mechanical stimulation on both the osteogenic and angiogenic capacity of osteocyte (OCY)-derived EVs [25, 26]; however, the impact of mechanical stimulation on the angiogenic potency of EVs derived from cells at earlier stages of the mesenchymal lineage remains unknown.

Mesenchymal stromal/stem cells (MSCs) have been shown to influence angiogenesis in a paracrine manner through both secreted soluble factors and EVs [27–31]. MSCs are sensitive to mechanical cues, and prior studies have explored how substrate stiffness can influence the angiogenic potential of their secretome [32]. Moreover, the secretome from osteogenically differentiated MSCs [10], as well as osteoblasts (OB) [33], have also been shown to promote angiogenesis, with murine osteoblasts releasing pro-angiogenic factors when subject to fluid flow [34]. However, the influence of mechanical stimulation on the angiogenic properties of the secretome and associated EVs from human MSCs and osteoblasts remains to be elucidated. The final stage of the lineage, osteocytes (OCY), can also influence endothelial cell behaviour [35] and mechanical stimulation enhances the angiogenic properties of both the secretome and EVs [26]. Yet, how the angiogenic properties of the mesenchymal bone cell changes with differentiation and whether mechanical stimulation can be used to tune these properties remains poorly understood.

Therefore, in this study we determined that mechanically stimulated MSCs, OBs and OCY derived secretomes and EVs had different effects on angiogenesis that was dependent on both the stage of lineage commitment and mechanical stimulation of the parent cell. Furthermore, as OCY-derived EVs proved to have the superior angiogenic efficacy, these EVs were chosen for further assessment, and were shown to elicit higher levels of endothelial cell migration and CD31 expression, indicating a more mature vessel phenotype. Moreover, these EVs were then assessed in a pre-clinical *ex ovo* chicken embryo chorioallantoic membrane (CAM) model, where superior vessel formation was once more seen with the mechanically-activated (MA) OCY-derived EV. This study thus delineates the role of mechanical stimulation and stage of lineage commitment on the angiogenic properties of bone cell derived EVs and importantly highlights a novel EV-based angiogenic therapy.

## Materials and Methods

### Cell culture

Human MSCs (hMSCs, Lonza) were maintained in DMEM high glucose GlutaMAX™ (Gibco™, 61965026) with 10% FBS (Foetal bovine serum) and 1% P/S.). GFP-expressing HUVECs (Human umbilical vein endothelial cells) (Lonza) were maintained in EBM-2 MC BulletKit medium (Lonza, CC-3156 and CC-4147). hMSCs in passages 3-5 were used for all experiments and HUVECs in passages 7-9 were used for angiogenesis experiments. Human osteoblasts (OBs) were obtained by treating hMSCs with osteogenic media (OM) (100 nM dexamethasone, 0.05 mM L-ascorbic acid and 10 mM β-glycerol phosphate) for 14 days, as detailed in the literature [25, 36, 37]. Serum starvation media for hMSCs and hOBs consisted of DMEM with 0.5% FBS, and 1% P/S. MLO-Y4 osteocyte cells (OCY) were maintained in α-MEM (Lonza, LZBE12-169F) with 2.5% FBS (Gibco, 10270106), 2.5% Calf Serum (CS) (Biosera, CA-115), and 1% P/S (Sigma Aldrich, P4333). Serum starvation media for MLO-Y4s consisted of α-MEM with 0.5% FBS, 0.5% CS and 1% P/S. MLO-Y4 (Kerafast) cells in passages 40-50 were used in all experiments.

### Mechanical stimulation and conditioned media collection

Cells were seeded on glass slides (Corning™, CLS294775X38) which were previously sterilised in 70% ethanol for 1 h, left to dry and coated with collagen type I (Corning™, 354236). MSCs/OBs were seeded at 0.61 × 10^4^ cells/cm^2^ and MLO-Y4 cells were seeded at 1.16 × 10^4^ cells/cm^2^. Following 24 h, the cells were treated with serum starvation medium to allow for cell cycle synchronisation. 48 h post seeding, the slides were placed in a custom-made parallel plate flow chamber bioreactor where oscillatory fluid shear was applied to the cells at 1 Pa and 1 Hz for 2 h [38]. The slides were then washed in PBS and placed in a custom-made slide chamber. 2.5 mL of serum starvation media with EV-depleted FBS and CS were added to the slides. Following 24 h in culture, the conditioned media (CM) was collected, centrifuged at 3,000 g for 10 min at 4°C and stored at −80°C until further use.

### Tube formation assay

To assess the effect of bone cell-derived CM, DM or EVs from bone cells on HUVEC angiogenesis, a Geltrex^TM^ (ThermoFischer, A1413302)-based tube formation assay was performed. Briefly, 0.1 ml of Geltrex^TM^ was added to each well of a 24-well plate. The coated plate was placed at 37°C for 30 min to allow for the matrix to solidify. Following complete gelation, 25,000 cells/cm^2^ were added to each well. CM or EVs were added to the cells at a 1:1 ratio with HUVEC growth media (with no added VEGF or serum). As a positive control, 10 ng/mL of VEGF (PeproTech®, 100-20) was added to the media. Following 18 h in culture, the plates were imaged using an Olympus IX83 inverted fluorescence microscope with 4x and 10x objectives. Quantification of the number of junctions, number of segments and total tube length was performed using the Angiogenesis Analyzer in ImageJ software (1.54).

### Extracellular vesicle collection

The protocol for EV collection was chosen in accordance with the ISEV minimal information guidelines [39]. EVs from MSCs, OBs and OCYs were collected from the collected CM by ultracentrifugation. Briefly, the CM was centrifuged at 10,000 g at 4°C for 30 minutes and filtered through a 0.45 μm pore filter. The supernatant was then transferred to ultracentrifugation tubes, which were weighed to ensure proper balance of the rotor (70Ti-fixed-angle). The tubes were centrifuged at 110,000 g for 75 min at 4°C using an Optima XPN-100 Ultracentrifuge. The pellets were then resuspended in PBS and centrifuged again in the same conditions. The final pellets were resuspended in 100 μL of filtered PBS. The leftover media without EVs was also stored as depleted media (DM).

### Extracellular vesicle characterisation

#### Transmission electron microscopy

10 µL samples of EV suspension were placed on formvar carbon-coated nickel grids (Ted Pella Inc, Cat.#:01813-F) and allowed settle for 10 min. A droplet of paraformaldehyde (2%) was placed on parafilm, and the grid was placed on top and fixed for 10 min. This was then contrasted in 2% uranyl acetate (BDH, Cat.#:230,550) and all images were taken using a JEOL JEM-2100 TEM (transmission electron microscopy) at 120 kV.

#### Nanoparticle tracking analysis

Nanoparticle tracking analysis (NTA) was conducted using the NTA NS300 system (NanoSight, Amesbury, UK). The NTA system allows for the determination of EV size distribution and concentration, measuring nanoparticles ranging from 10 nm to 1000 nm. This system utilizes both light scattering and Brownian motion. When nanoparticles encounter laser beams, they scatter light, which is then visualized at a 20X magnification. Brownian motion of the particles is observed by capturing 30 frames per second. Filtered PBS (0.45 μm filter) served as a negative control. EV samples were appropriately diluted with filtered PBS, loaded onto the NTA using a NanoSight syringe pump, and subjected to five 60-second video recordings. The NTA software was then used to determine the size of the particles.

#### Flow cytometry

EV surface antigens were exposed to antibodies diluted in 0.22 μm-filtered PBS with 2% EV-depleted FBS supplemented with protease inhibitor and phosphatase inhibitor (IFCM buffer). The antibodies used were anti-CD63 conjugated with FITC (1:15) (Biolegend, Cat. #: 353006), and CD81-PE-Cy7 (1:15) (Biolegend, Cat. #: 349512). The EVs were incubated with the antibodies for 45 min at room temperature in the dark, and washed using a 300 kDa filter (Nanosep, Cat. #: 516-8531), resuspended in 50 μL IFCM buffer and acquired within 2 h on the ImageStream X MK II imaging flow cytometer (Amnis/Luminex, Seattle, USA) at 60x magnification and low flow rate. EV-free IFCM buffer and unstained EVs were run in parallel. Fluorescence was within detection linear range in the following channels: FITC was measured in channel 2 (B/YG_480-560 nm), and PE-Cy7 in channel 6 (B/YG_ 745-780 nm). Brightfield in channel 1 and 9 (B/YG_435-480 and R/V_560-595 nm filter, respectively) and side scatter channel (SSC) in channel 12 (R/V_745-780 nm 49 filter). Data analysis performed using IDEAS software v6.2 (Amnis/Luminex, Seattle, USA). EVs were gated as SCC-low vs fluorescence, then as non-detectable brightfield (Fluorescence vs Raw Max Pixel Brightfield channel), gated EVs were confirmed in IDEAS Image Gallery.

#### Protein concentration

EV samples derived from both mechanically stimulated and statically cultured cells were lysed by cell lysis buffer (Thermo Fisher Scientific, Cat. #: FNN0011) supplemented with protease inhibitor cocktail (Roche, Basel, Switzerland; Cat. #: 04693116001). The samples were then vortexed for 10 s and placed on ice, with this process repeated 3 times. Following lysis, samples were then centrifuged at 13,200 g for 10 min at 4°C and the supernatant collected for analysis. The protein content in the EV lysate was quantified using Pierce™ BCA Protein Assay Kit (Thermo Scientific™, 23225) according to manufacturer’s instructions.

### Proliferation assay

The assay was performed as per manufacturer’s descriptions (Roche, 11647229001). Briefly, HUVECs were seeded at 12,500 cells/cm^2^ and treated with collected EVs, or with VEGF (10 ng/ml) as a positive control. Following 24 h, BrDU was added to each sample and incubated for a further 24 h. Anti-BrDU solution was added, and the samples were analysed through a plate reader at 450 nm.

### Migration assay

The Transwell assay was performed in which hanging cell culture inserts (PIEP12R48, Merck Millipore) were placed into well plates with serum-free media in each well. HUVECs were resuspended in serum free media and 10,000 cells were added to the upper side of the of the 8 µm pore hanging cell culture inserts. Cells were incubated in serum free media for 1-4h to allow adhesion. The inserts were then transferred to wells containing the EVs or VEGF (10 ng/mL) and 10% FBS as positive controls. The plate was incubated for 18 h and cells were fixed in neutral buffered formalin (NBF) and stained with pure hematoxylin (Sigma HHS32-1L). Following an 8 min of incubation, wells were rinsed 3 times with diH_2_O. Non-migrated cells were wiped from the membrane and the membrane was mounted on a glass slide. Slides were imaged using a 20x lens and cell number was counted in 8 fields of view.

### Immunocytochemistry

Following the same protocol as for tube formation, 18 h after incubation on Geltrex^TM^, cells were fixed with 10% formalin for 10 min, followed by washes in PBS. Subsequently, cell membranes were permeabilised using 0.2% Triton X-100 for 10 min and blocked with 3% BSA for 30 min at room temperature (RT). The cells were incubated with the primary antibody (CD31+, 1:200 dilution) overnight at 4°C, followed by incubation with the second antibody (Alexa 488, 1:1000 dilution) for 1 h at RT. Finally, the cells were incubated with DAPI (1:1000 dilution) for 10 min and washed prior to imaging using an Olympus IX83 inverted fluorescence microscope with a 10x objective. Mean fluorescence intensity was determined using ImageJ.

### Ex ovo chick chorioallantoic membrane assay

The chorioallantoic membrane (CAM) assay is an established *ex ovo*, shell-less chicken embryo model, to determine the angiogenic potential of tissue engineered therapies. All experimentation carried out on chick embryos was in accordance with the EU Directive 2010/63/EU for animal experiments. Fertilised chicken eggs (Shannonvale foods, Co. Cork, Ireland) were incubated horizontally at 37.5 °C with 60% of humidity for 70-72 h. On day 3, the eggs content were placed into 100×20 mm petri dishes (Corning Inc., New York, USA). On day 7 of development, 6 mm filter paper discs containing EVs resuspended in PBS (PBS only as a control) were placed on the membrane and incubated for 5 additional days, reaching a total of 10 days in culture. To assess the local effect of the scaffolds on the angiogenesis of the chick embryo model, microscopy images were taken and assessed using a FIJI macro [40] which utilised a MorphoLibJ[41]. The analysed vessels yielded the number of branches, junctions, vessel area and maximum branch length for each processed field of view.

### Statistical analysis

All results are expressed as mean ± standard deviation (SD) and statistical analysis was performed using GraphPad Prism 10.0 software. All data is assessed for normality with the Shapiro-Wilk test. Significant differences reported when tested using a one-way analysis of variance (ANOVA) with Tukey’s Multiple Comparison Test or a Kruskal–Wallis test, depending on the normality of the data. Significance is represented as *p≤ 0.05, **p≤ 0.005, ***p≤ 0.001, **** p≤0.0001. Number of experimental replicates are reported as n.

**Scheme 1.**
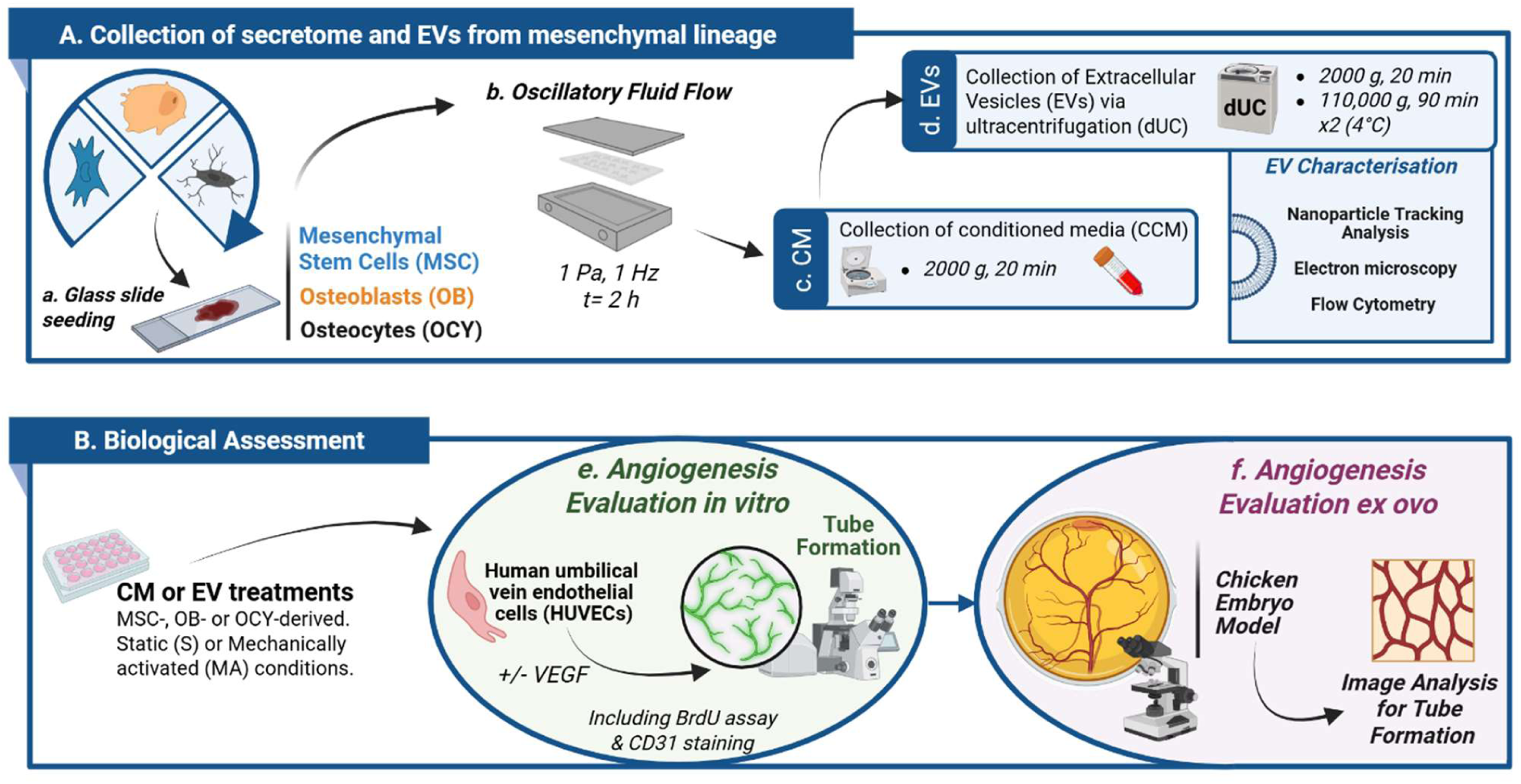
Schematic of experimental workflow for mechanically activated conditioned media and EV collection with angiogenic assessment. **(A)** Collection of secretome and EVs from mesenchymal lineage **(B)** Biological assessment of CM and EVs. Created using Biorender.com

## Results

### The efect of the mechanically stimulated MSC secretome on the tube formation of HUVECs

To determine the angiogenic properties of paracrine factors released by both static- and mechanically-activated hMSCs, HUVECs were treated with conditioned media (CM) collected from hMSCs and a tube formation assay was performed. Fluorescent images of GFP-tagged HUVECs presented with evident mesh and node formation in the positive control group (+VEGF) when compared to the negative control group (-VEGF) (Fig. 1A), an effect which was verified upon quantification of number of junctions, segments, or total tube length (p<0.05, Fig. 1B-D) demonstrating successful angiogenesis. However, upon treatment with MSC derived static conditioned media (MSC-CM^S^) there was no apparent change in angiogenesis when compared to the negative control (Fig. 1A-D), indicating that MSCs do not secrete factors which influence angiogenesis under static culture conditions utilised in this study. On the contrary, visible junctions and segments are evident in HUVECs treated with the mechanically-activated MSC-derived conditioned media group (MSC-CM^MA^). This was confirmed through quantification, where there was a significant increase in both the number of junctions and number of segments when compared to the −VEGF group (Fig. 1B-D). This increase led to tube formation values on par with the positive control (+VEGF). Furthermore, treatment with MSC-CM^MA^ lead to a trend of increased total tube length, with a 1.4-fold increase compared to −VEGF. Therefore, paracrine factors released by mechanically-activated MSCs demonstrate a favourable angiogenic effect.

**Figure 1.**
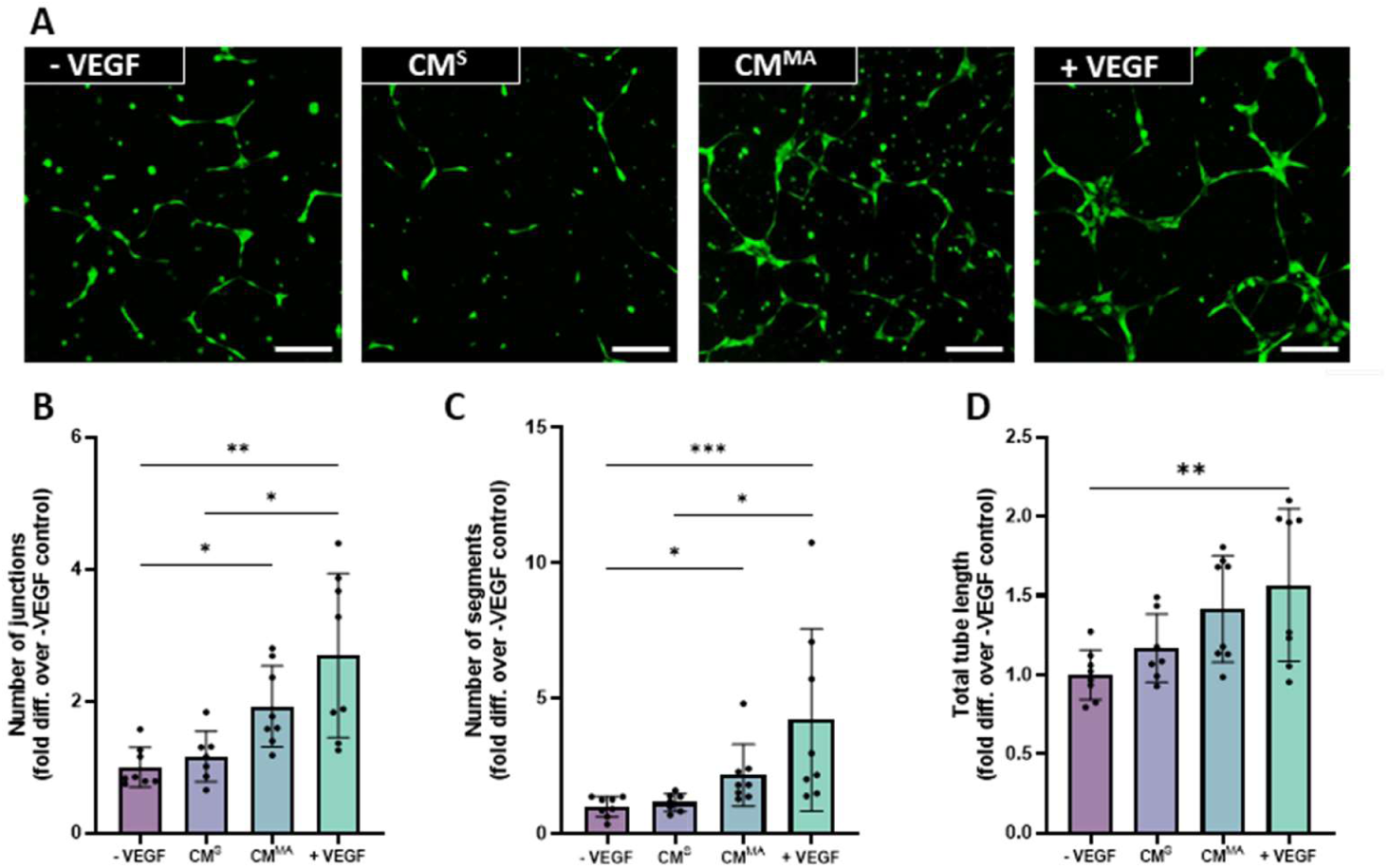
HUVEC angiogenesis in response to hMSC-derived static and mechanically activated conditioned media. **(A)** Fluorescent images of tube formation at 18 h. **(B)** Number of junctions **(C)** Number of segments and **(D)** Total tube length normalised to negative control. Scale bar = 200 µm. Significant differences reported when tested using a Kruskal–Wallis test **(B, C)** or one-way ANOVA **(D)** Data presented as Mean ± SD, n ≥ 6.

### The efect of the mechanically stimulated OB secretome on the tube formation of HUVECs

To determine how the stage of lineage commitment affects the angiogenic properties of the bone cell secretome, MSCs were differentiated into OBs through osteogenic supplementation for 14 days (Fig. 2A). Differentiation was confirmed by an increase in ALP activity when compared to the BM (basal media) treated MSCs (Fig. 2B) and positive calcium staining (Fig. 2C, D). Upon osteoblastic differentiation, OBs were lifted from the mineralised matrix through a collagenase and trypsin treatment (Fig 2E) and reseeded and placed in the PPFC (parallel plate flow chamber) bioreactor. OBs were then exposed to mechanical stimulation and CM was collected and utilised to treat HUVECs as above. Similarly to MSC-CM^S^, CM collected from statically cultured osteoblasts (OB-CM^S^) did not influence angiogenesis (Fig. 2F-I) when compared to both the −VEGF control. Interestingly, CM collected from mechanically activated osteoblasts (OB-CM^MA^) did not elicit a significant angiogenic response (Fig. 2F) despite a small increase, as measured by number of junctions, segments and total tube length when compared to −VEGF control (Fig. 2F-I). This is in contrast to MSC-CM^MA^ indicating a shift in the angiogenic properties of the mesenchymal bone cell secretome with progression along the osteogenic lineage.

**Figure 2.**
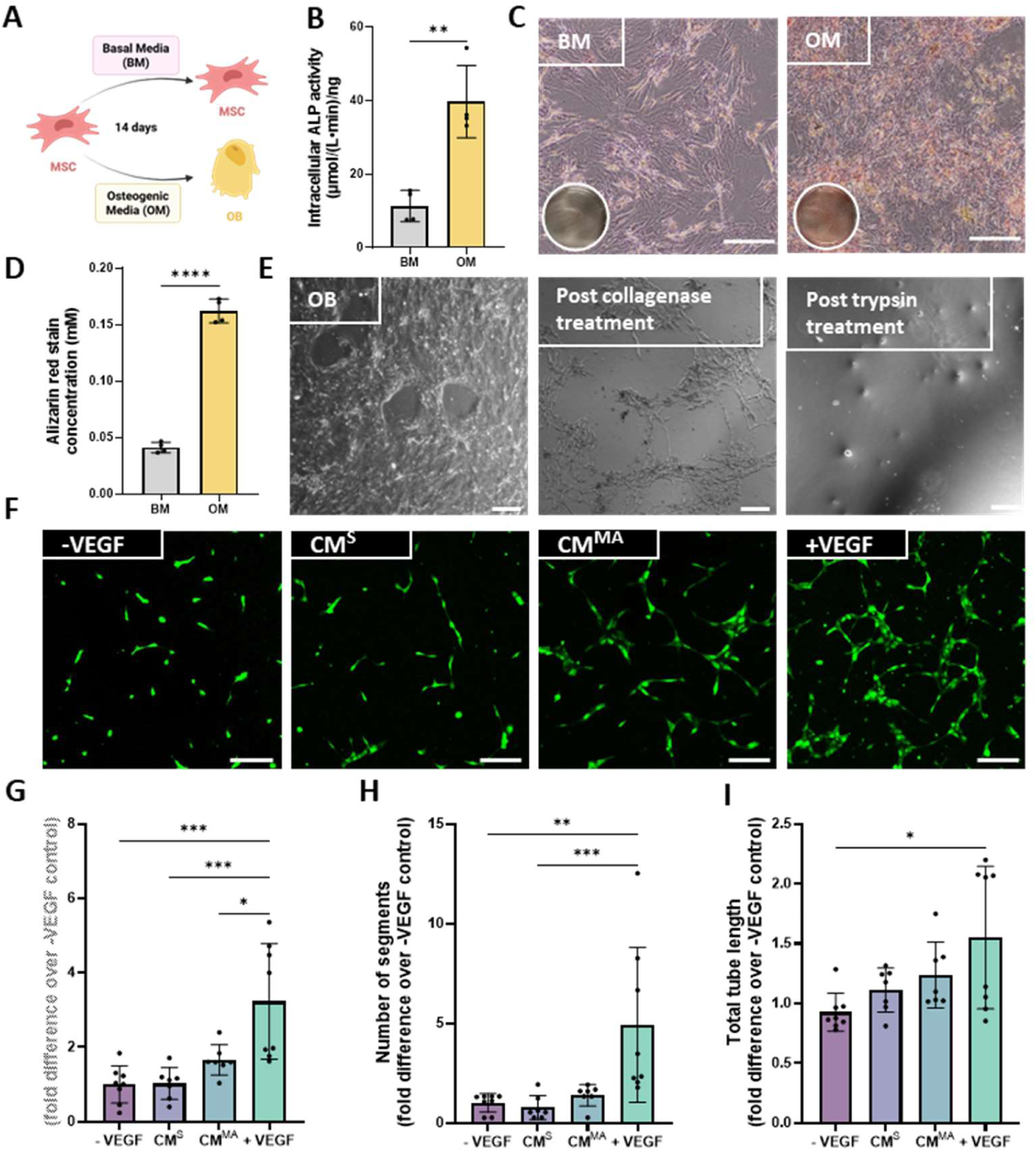
HUVEC angiogenesis in response to hOB-derived static and mechanically activated conditioned media. **(A)** Schematic of osteoblast differentiation experimental design**. (B)** Intracellular ALP activity at 14 days normalised to DNA **(C)** Alizarin red staining of control and osteoblast at day 14 **(D**) Quantification of alizarin red staining **(E)** Protocol for trypsinisation of hOBs. **(F)** Fluorescence microscopy images of tube formation at 18h. **(C)** Number of junctions **(D)** Number of segments and **(E)** Total tube length normalised to negative control. Scale bar= 200 µm. Significant differences reported when tested using an unpaired t-test **(B, D),** one-way ANOVA **(G)** or a Kruskal–Wallis test **(H,I).** Data presented as Mean ± SD, n≥6.

### The efect of the mechanically stimulated OCY secretome on the tube formation of HUVECs

To further assess the impact of the stage of lineage commitment on the angiogenic potential of the bone cell secretome, both static- and mechanically-activated-CM derived from the final stage of differentiation, osteocytes, were used to treat HUVECs. As seen with MSC-CM^S^ and OB-CM^S^, CM collected from statically-cultured osteocyte (OCY-CM^S^) did not influence angiogenesis (Fig. 3A), further verifying that mesenchymal-derived bone cells do not secrete factors which influence angiogenesis under static culture conditions utilised in this study. With regards to the angiogenic properties of paracrine factors released by mechanically activated osteocytes (OCY-CM^MA^), distinct junctions and segments can be seen following OCY-CM^MA^ treatment (Fig. 3A), however upon quantification and despite a 1.3- and 1.4-fold increase in number of junctions and segments when compared to −VEGF control, these values did not reach significance (Fig. 3B-D).

**Figure 3.**
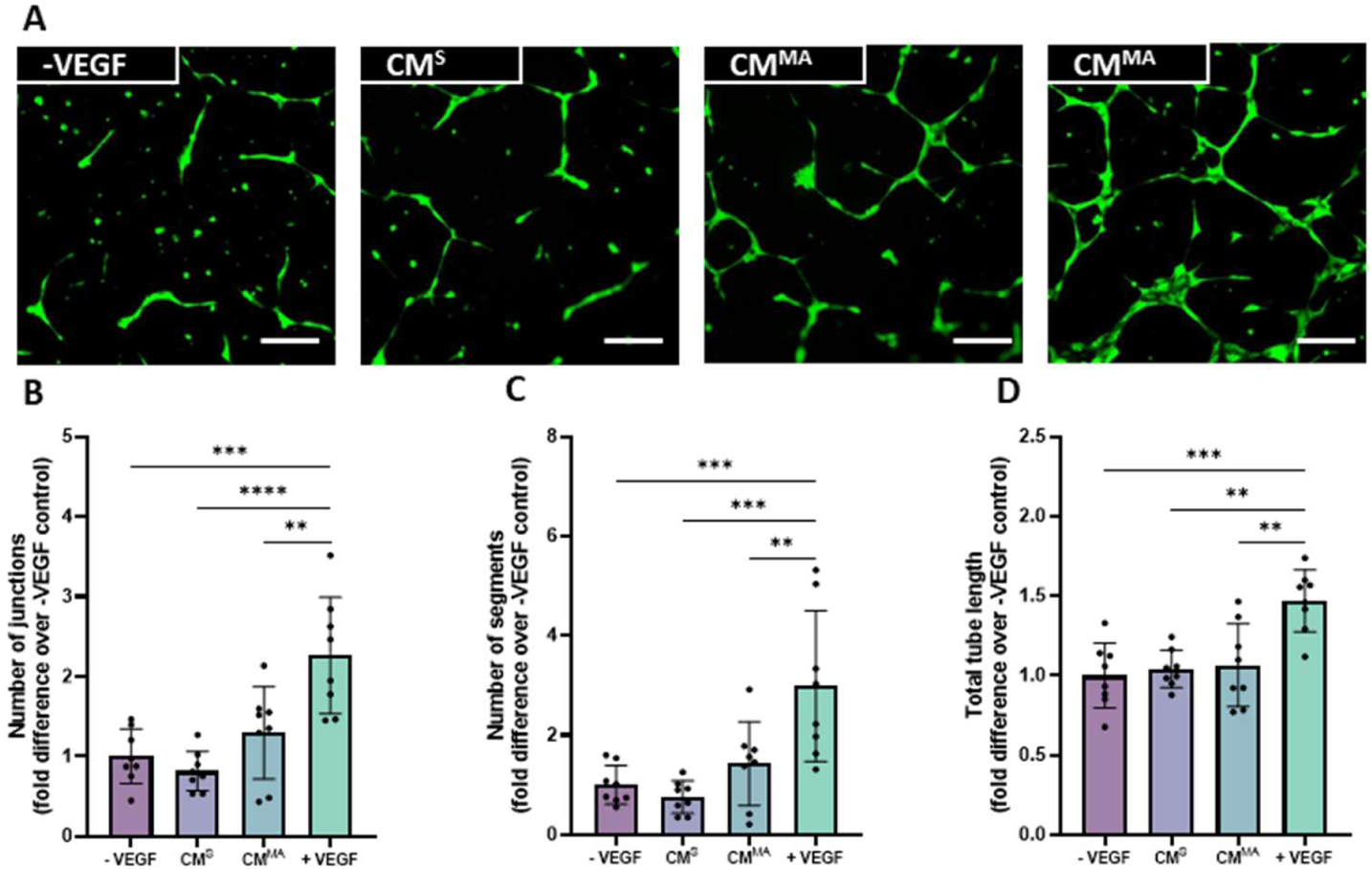
HUVEC angiogenesis in response to OCY-derived static and mechanically activated conditioned media. **(A)** Fluorescent images of tube formation at 18hrs. **(B)** Number of junctions **(C)** Number of segments and **(D)** Total tube length normalised to negative control. Scale bar= 200 µm Significant differences reported when tested using a one-way ANOVA. Data presented as Mean ± SD, n=8.

Taken together, these results indicate that the angiogenic properties of the mesenchymal bone cell secretome is lost with differentiation along the osteogenic lineage and that these properties can be enhanced by mechanical stimulation of the parent cell.

### EVs released from mesenchymal-derived bone cells display similar characteristics along the osteogenic lineage

EVs have emerged as key mediators of intercellular communication that can influence various physiological processes, including angiogenesis. To investigate whether EVs present in the secretome were responsible for the angiogenic effects demonstrated with the whole secretome, EVs were collected via ultracentrifugation from CM obtained from MSCs, OBs and OCYs. EVs derived from all cell types exhibited the typical round shaped morphology, as shown by TEM images (Fig. 4A). Moreover, all EVs showed an average size of approximately 140 nm (Fig. 4B, C). Interestingly, OCY-EVs showed a trend of higher particle concentration as determined by NTA compared to MSC-EVs and OB-EVs (Fig. 4D). However, similar EV yield was obtained across the mesenchymal lineage with no significant differences in EV concentration observed. Positive staining for tetraspanin markers CD63 and CD81 through flow cytometry (Fig. 4E), further confirming the successful collection of EVs. The protein content per volume of collected conditioned media was approximately 0.5 µg/mL for all cell types (Fig. 4F). Therefore, a treatment of 0.5µg/mL was chosen for further angiogenic assessment, so as to correspond to the same concentration present in the CM.

**Figure 4.**
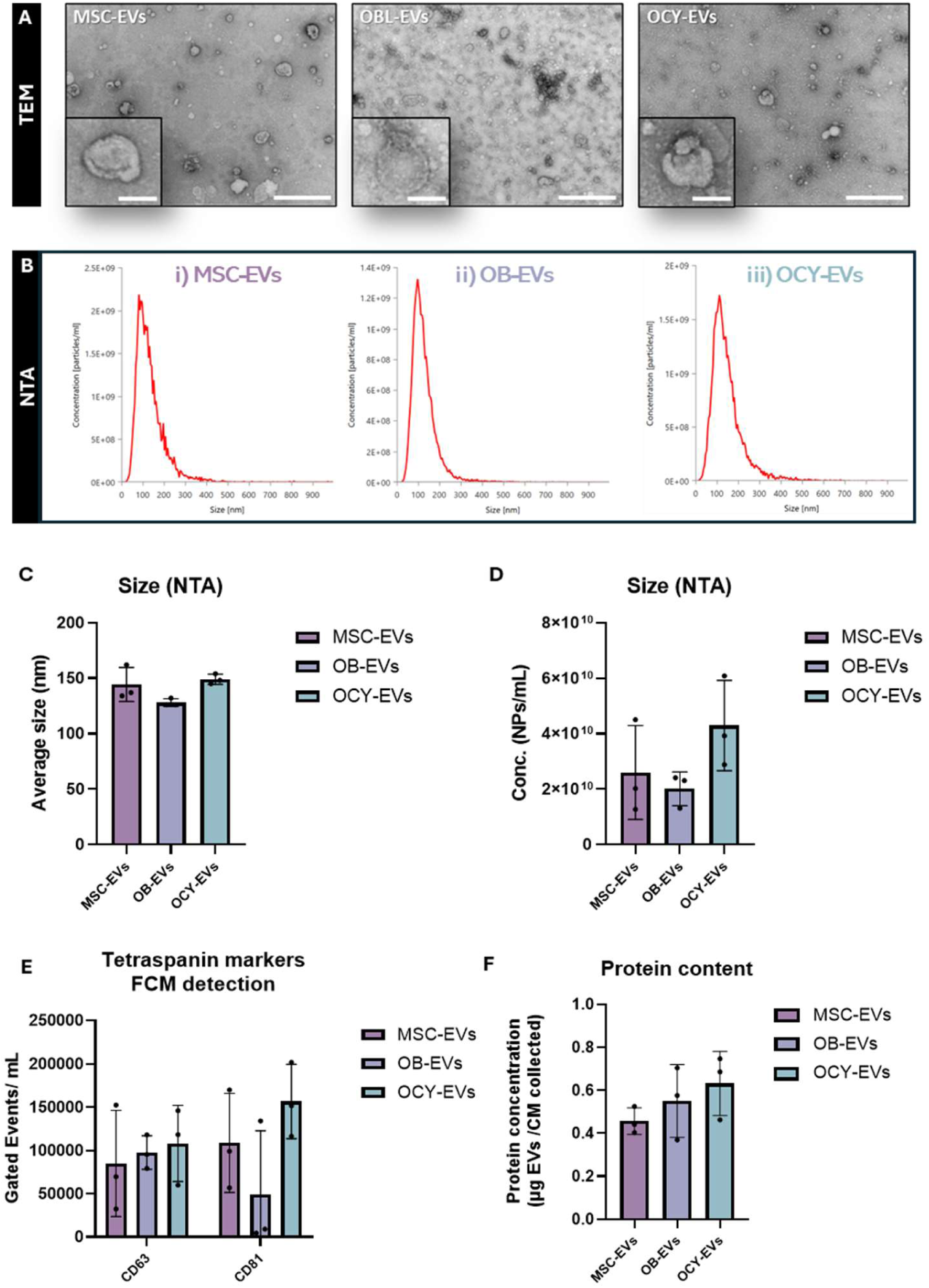
Comparative collection and characterisation of EVs derived from the conditioned media of MSCs, OBs and OCYs. **(A)** TEM images of EVs, scale bars = 500 nm (100nm – inserts). NTA was performed to determine **(B)** Size Distributions **(C)** Average Nanoparticle Sizes and **(D)** Nanoparticle Concentrations. **(E)** Detection of tetraspanin markers CD63 and CD81 via flow cytometry. **(F)** Protein content via the BCA assay. Data presented as Mean ± SD, n=3.

### MSC-derived EVs do not influence angiogenesis

To determine whether EVs were responsible for the angiogenic effects seen with the whole secretome, both the collected EVs and the conditioned media depleted of these EVs (DM) were used to treat HUVECs, and tube formation was once more assessed. Treatment with EVs collected from statically cultured MSCs (MSC-EV^S^) demonstrated a slight decrease in junctions and segments when compared to the negative control although this was not found significant. A similar trend was seen in the static depleted media group (DM^S^) (Fig. 5A), which was confirmed through quantification (Fig. 5B, C), and is consistent with treatments with the whole secretome as determined previously (Fig. 1). Interestingly, mechanical stimulation enhances the angiogenic properties of the collected EVs although no differences were seen in the number of junctions, segments and total tube length when compared to the negative control (Fig. 5A-D). The mechanically activated EV-depleted media (DM^MA^) showed a slight increase in all three readouts compared to the EVs (Fig. 5B-D), which could indicate that a combination of soluble factors and EVs are responsible for the increase in tube formation as observed with the whole CM. Nonetheless, similar results to that of junctions and segments can be seen with total tube length across all groups (Fig. 5D).

**Figure 5.**
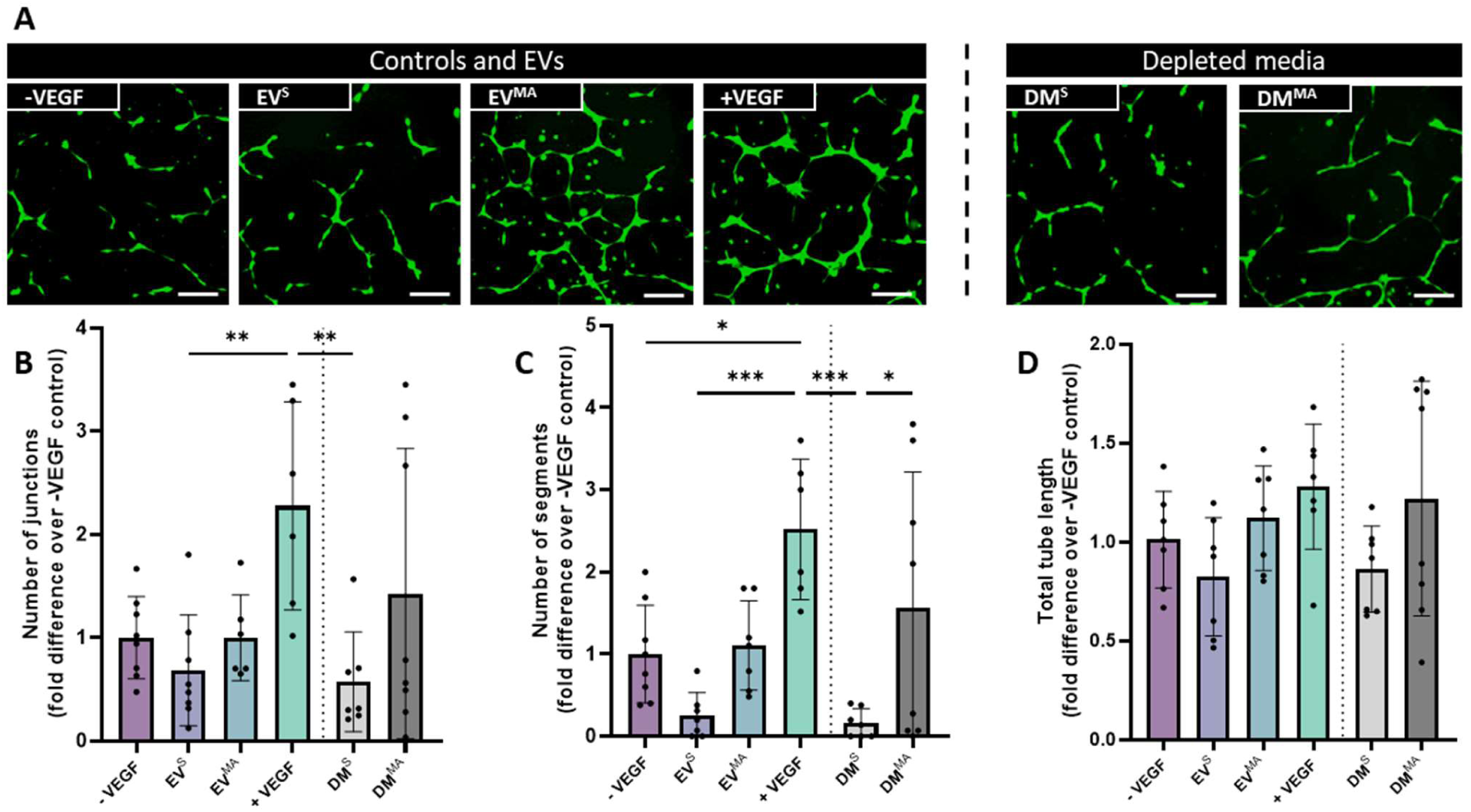
HUVEC angiogenesis in response to MSC-derived static EVs, mechanically activated EVs, and EV depleted media. **(A)** Fluorescent images of tube formation at 18hrs. **(B)** Number of junctions **(C)** Number of segments and **(D)** Total tube length normalised to negative control. Scale bar= 200 µm Significant differences reported when tested using a one-way ANOVA. Data presented as Mean ± SD, n≥6.

### OB derived EVs enhance tube formation of HUVECs

EVs derived from OBs were then used to treat HUVECs, to assess the effect of the stage of lineage on the EV angiogenic potency (Fig. 6A). OB-EV^S^ appear to induce a slight increase in number of junctions and segments, although not significant, which is on par with the static depleted media (Fig. 6A-C) and is consistent with that seen following treatment with the whole secretome (Fig. 2). However, the OB-EV^MA^ groups show increased number of junctions and segments, on par with the positive control (Fig. 6B, C), indicating a positive effect of mechanical stimulation on the angiogenic potency of these EVs that was not seen in CM^MA^ only. Moreover, the DM^MA^ has a slight trend of decreased number of junctions and segments compared to the EV^MA^, which may explain the lack of effect observed previously with the whole secretome. Similarly to the MSC-EVs, the total tube length follows similar trends to that of junctions and segments for all groups (Fig. 6D). Altogether these results show that differentiating MSCs into osteoblasts leads to the release of EVs that have a pro-angiogenic effect when these cells have been mechanically stimulated. This result highlights the importance of the stage of lineage on the regenerative effect of EVs.

**Figure 6.**
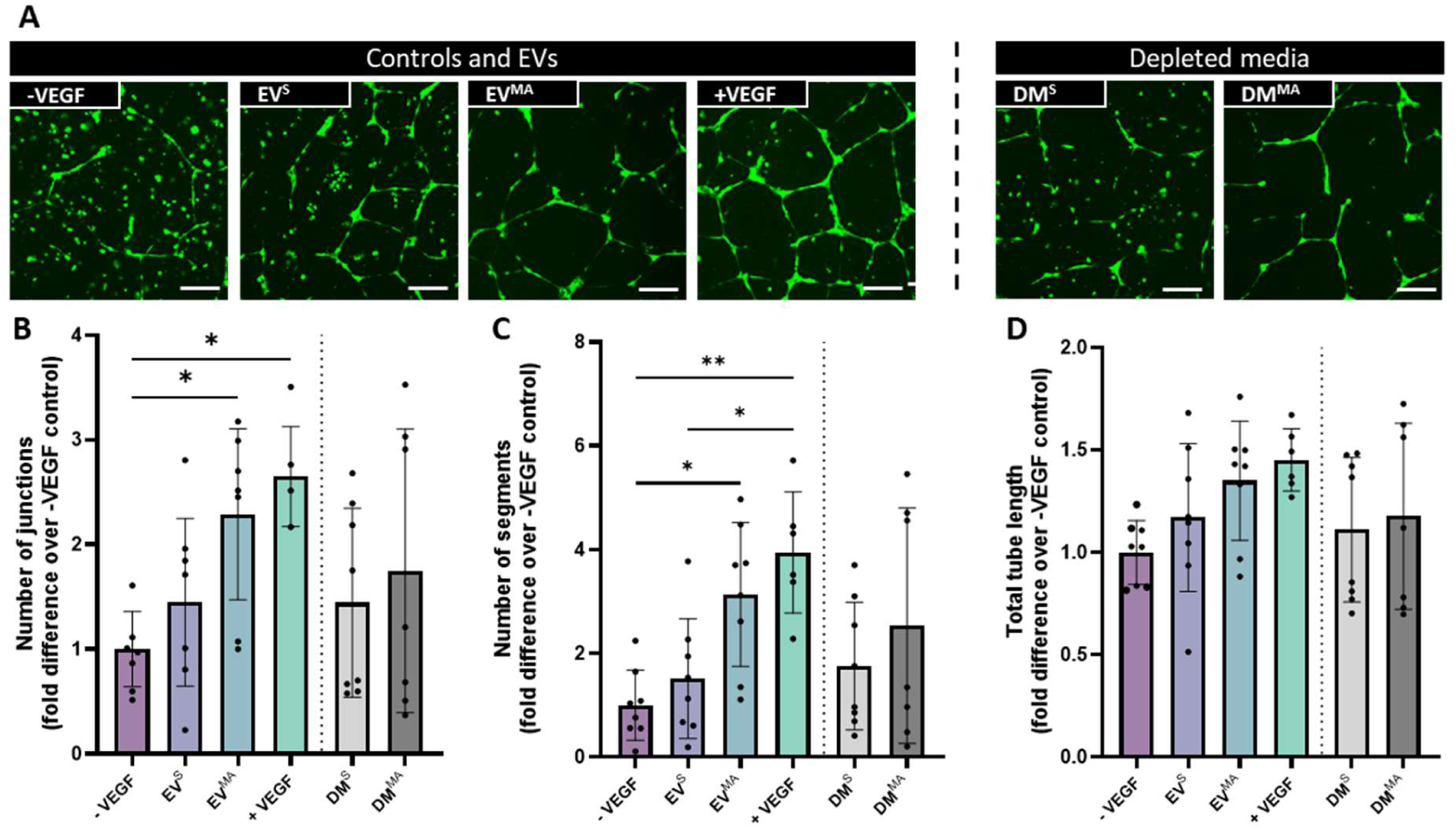
HUVEC angiogenesis in response to OB-derived static EVs, mechanically activated EVs, and EV depleted conditioned medias. (A) Fluorescent images of tube formation at 18hrs. **(B)** Number of junctions **(C)** Number of segments and **(D)** Total tube length normalised to negative control. Scale bar= 200 µm Significant differences reported when tested using a one-way ANOVA. Data presented as Mean ± SD, n≥6.

### OCY derived EVs enhance tube formation of HUVECs

Finally, the angiogenic potency of EVs derived from the terminally differentiated osteocytes was assessed (Fig. 7A). EV^s^ had no effect on junctions, segments or total tube length, on par with both the DM^S^ and CM^S^ (Fig. 7A-D). On the contrary, OCY-EVs^MA^ show clear junction and node formation (Fig. 7A), which are confirmed with a significant increase in all three readouts, on par with the +VEGF control (Fig.7B-D). Furthermore, mechanically-activated CM depleted of EVs (DM^MA^) has an inhibitory effect compared to the EV^MA^s and +VEGF control, as well as a trend of decreased tube formation when compared to the −VEGF control (Fig. 7A-D). This could indicate the presence of anti-angiogenic soluble factors in the media, which in turn explain the lack of angiogenic response in the full conditioned media groups.

**Figure 7.**
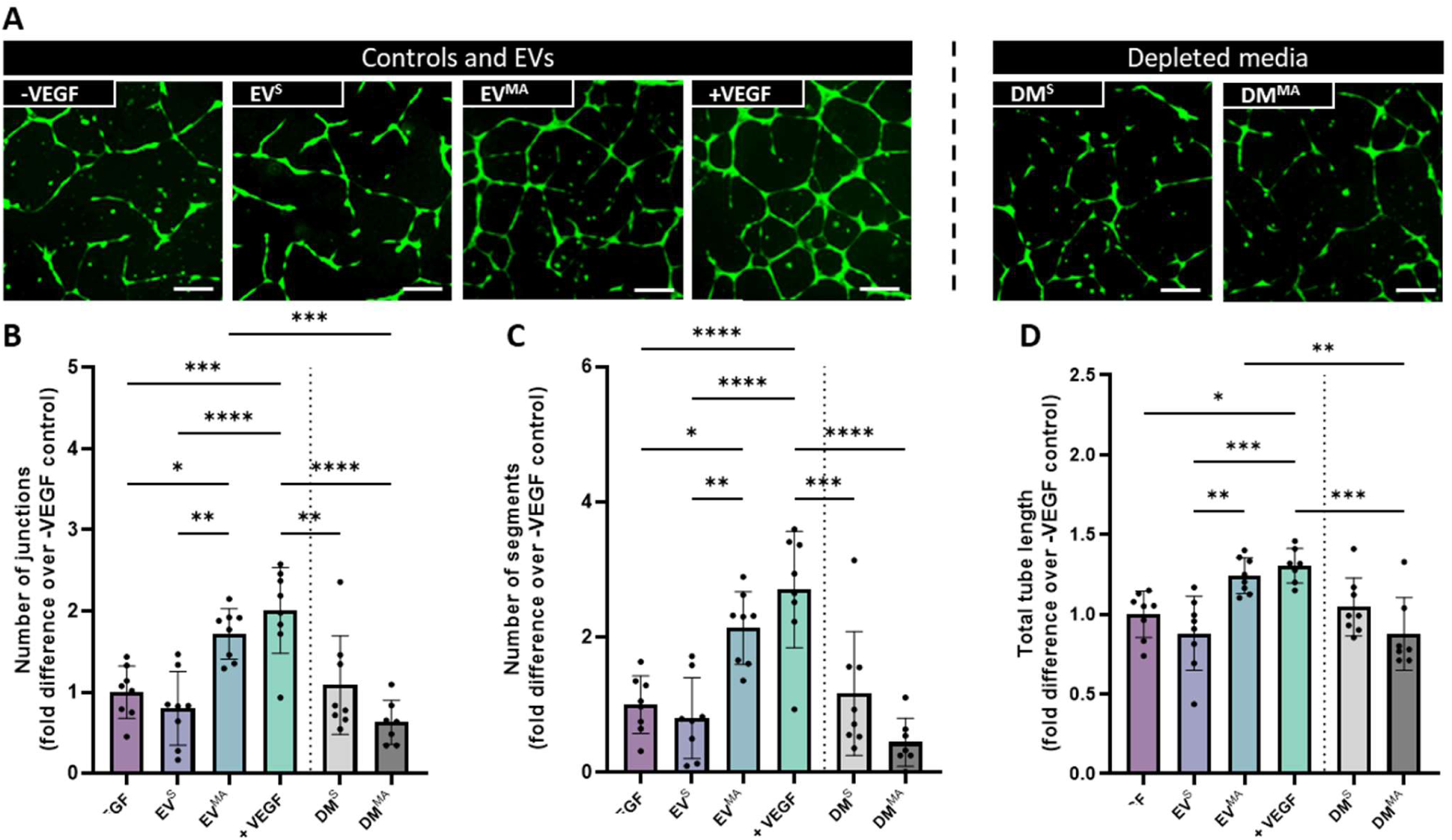
HUVEC angiogenesis in response to OCY-derived static EVs, mechanically activated EVs, and EV-depleted conditioned media. **(A)** Fluorescent images of tube formation at 18h. **(B)** Number of junctions **(C)** Number of segments and **(D)** Total tube length normalised to negative control. Scale bar= 200 µm. Significant differences reported when tested using a one-way ANOVA. Data presented as Mean ± SD,n≥6.

Interestingly, EVs^MA^ are the only EV type where a significant increase in total tube length can be seen when compared to the −VEGF control (Fig. 7D). These results further highlight the impact of lineage and the importance of mechanical stimulation on the angiogenic properties of secreted EVs and underscore osteocyte-derived EVs as a possible angiogenic therapy. Therefore, OCY-EVs^MA^ were chosen for further advanced characterisation as they elicited the optimal angiogenic response.

### The angiogenic response of OCY-EVs^MA^ is inversely correlated to EV dose

Since it was observed that OCY-EV^MA^ had the most robust response between the three cell types assessed, the optimal dose was investigated. To determine this optimal dose, three different doses of OCY-derived EVs (0.5 µg/mL, 2 µg/mL and 4 µg/mL) were used to treat HUVECs, and their tube formation was assessed (Fig. 8A). Focusing on EV^S^, increasing the dose beyond 0.5 µg/mL (concentration present in CM) did not improve the angiogenic response, and a slight decrease in the number of junctions and segments was identified at 2 µg/mL when compared to the negative control, although this was not significant. Similarly, with the EV^MA^ groups, there is a consistent effect with all the concentrations; however, as the concentration increases the variation between samples increases, rendering higher doses less reliable to use as treatments (Fig. 8B). Moreover, there is a dose-dependent decrease in the number of segments as the concentration of EVs^MA^ increases (Fig. 8C). Since EVs^MA^ exhibit the most favourable results in terms of junctions and segments at 0.5 µg/mL, while also demonstrating lower variability compared to other doses, this concentration was selected as the optimal dose for further studies.

**Figure 8.**
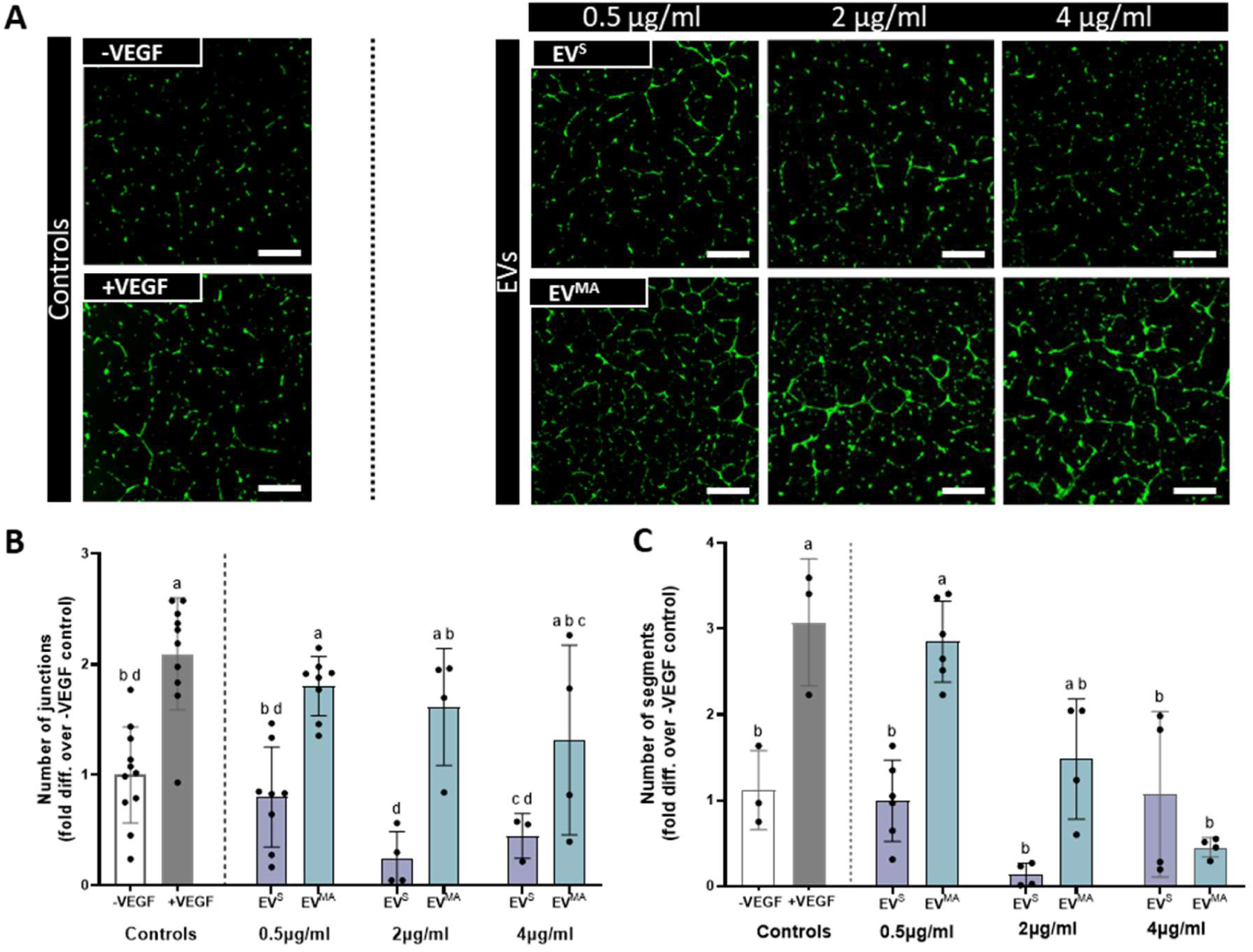
Dose response of osteocyte-derived static and mechanically activated EVs on HUVEC tube formation. **(A)** Fluorescence microscopy images of tube formation at 18h. **(B)** Number of junctions **(C)** Number of segments normalised to negative control. Scale bar= 200 µm. Significant differences reported when tested using a one-way ANOVA. Data presented as Mean ± SD, n≥3, N=2. Significant differences are represented according to the compact letter display (CLD).

### Treatment with OCY-EV^MA^ leads to increased endothelial cell migration and CD31 expression in HUVECs

Angiogenesis is a complex process that involves the coordinated proliferation and migration of endothelial cells to form new blood vessels. Therefore, in order to have a more advanced characterisation of this process, the effect of OCY-derived EVs on HUVEC proliferation and migration was assessed through BrdU incorporation and a transwell assay respectively. Although there were no significant differences between the EV groups and the controls in terms of cell proliferation (Fig. 9A), there was a significant increase in cell migration when HUVECs were treated with EV^MA^ when compared to the negative control and EV^s^ (p<0.05, Fig. 9B). Additionally, the expression of CD31, a marker of mature vessel formation, was assessed through immunofluorescence staining (Fig. 9C). Remarkably, there was a notable increase in expression of this marker when HUVECs were treated with EV^MA^ compared to all groups, including the +VEGF control (Fig. 9D). Therefore, OCY-derived EV^MA^ can improve the migration of HUVECs and form more mature tubules that highly express CD31, demonstrating the impact of mechanical conditioning of the parent cell in enhancing the angiogenic properties of the released EVs.

**Figure 9.**
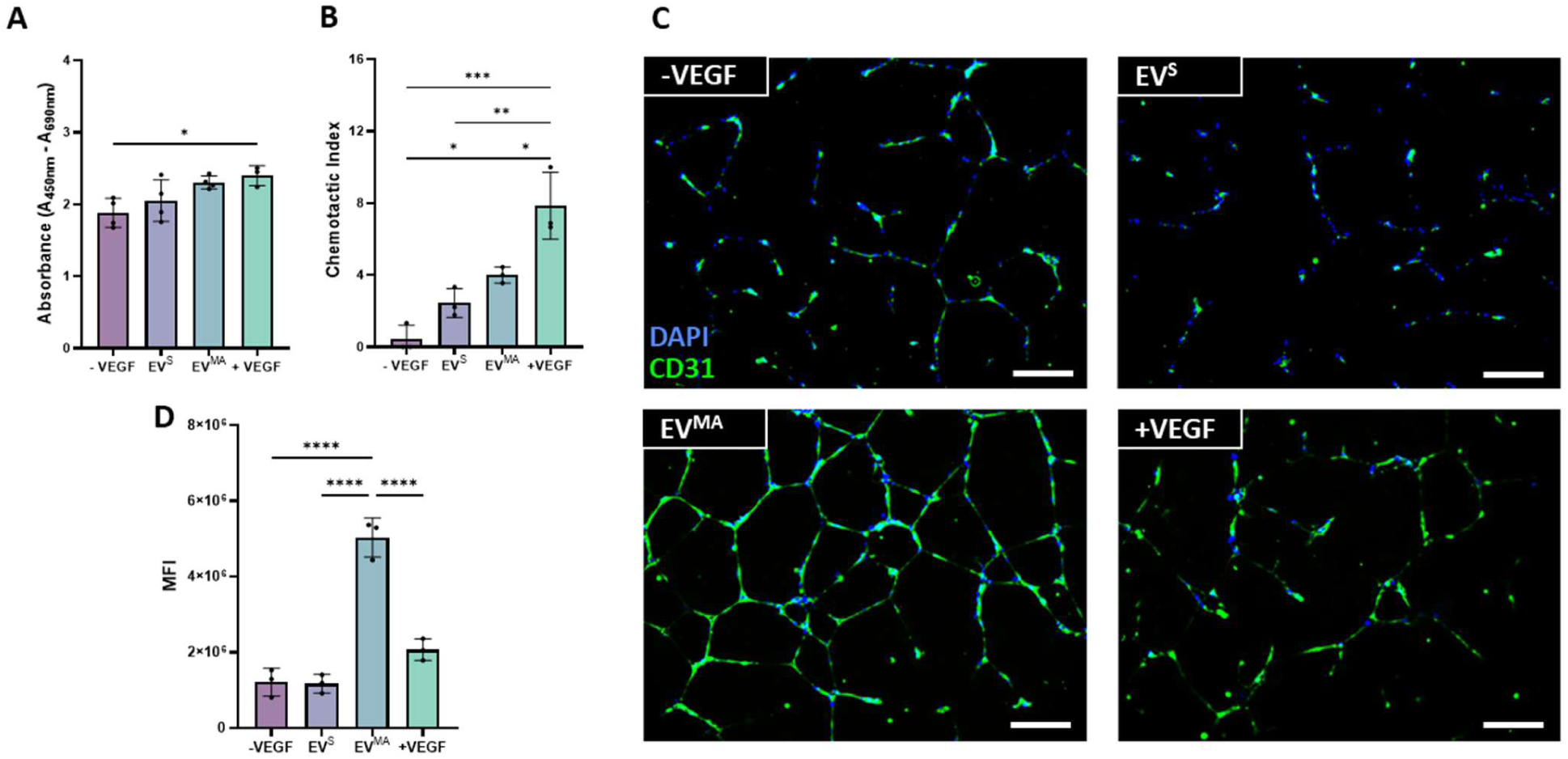
Advanced characterization of osteocyte-derived EVs on angiogenesis. **(A)** BrdU incorporation absorbance **(B)** Chemotactic index of static and mechanically activated EVs **(C)** CD31+ and DAPI immunofluorescent staining **(D)** Mean intensity quantification of CD31+ staining. Significant differences reported when tested using a one-way ANOVA. Data presented as Mean ± SD, n=3.

### Treatment with OCY-EV^MA^ leads to increased angiogenesis in an ex ovo chick embryo model

To assess the effect of these EVs in a more physiologically relevant model, an *ex-ovo* chick embryo CAM assay was performed. 0.5 µg/mL of EVs were loaded onto filter paper discs and positioned over sprouting blood vessels in the CAM at day 7 of incubation. Following a further 5 days in culture, images were taken to assess the vessel formation within the CAMs of the embryos (Fig. 10A). Vessel formation was quantified around each EV-disc (Fig. 10B). Similarly to the *in vitro* tube formation assay, OCY-EV^s^ did not affect vessel formation in this more physiologically-relevant model. Although following treatment with OCY-EV^MA^ there is a trend of increased number of junctions and segments this effect did not reach significance. However, the treatment with EV^MA^ significantly increased the total tube length when compared to control. These results are consistent with *in vitro* tube formation results, where OCY-EVs^MA^ demonstrated increases in HUVEC total tube length further highlighting the possible therapeutic potency of these EVs.

**Figure 10.**
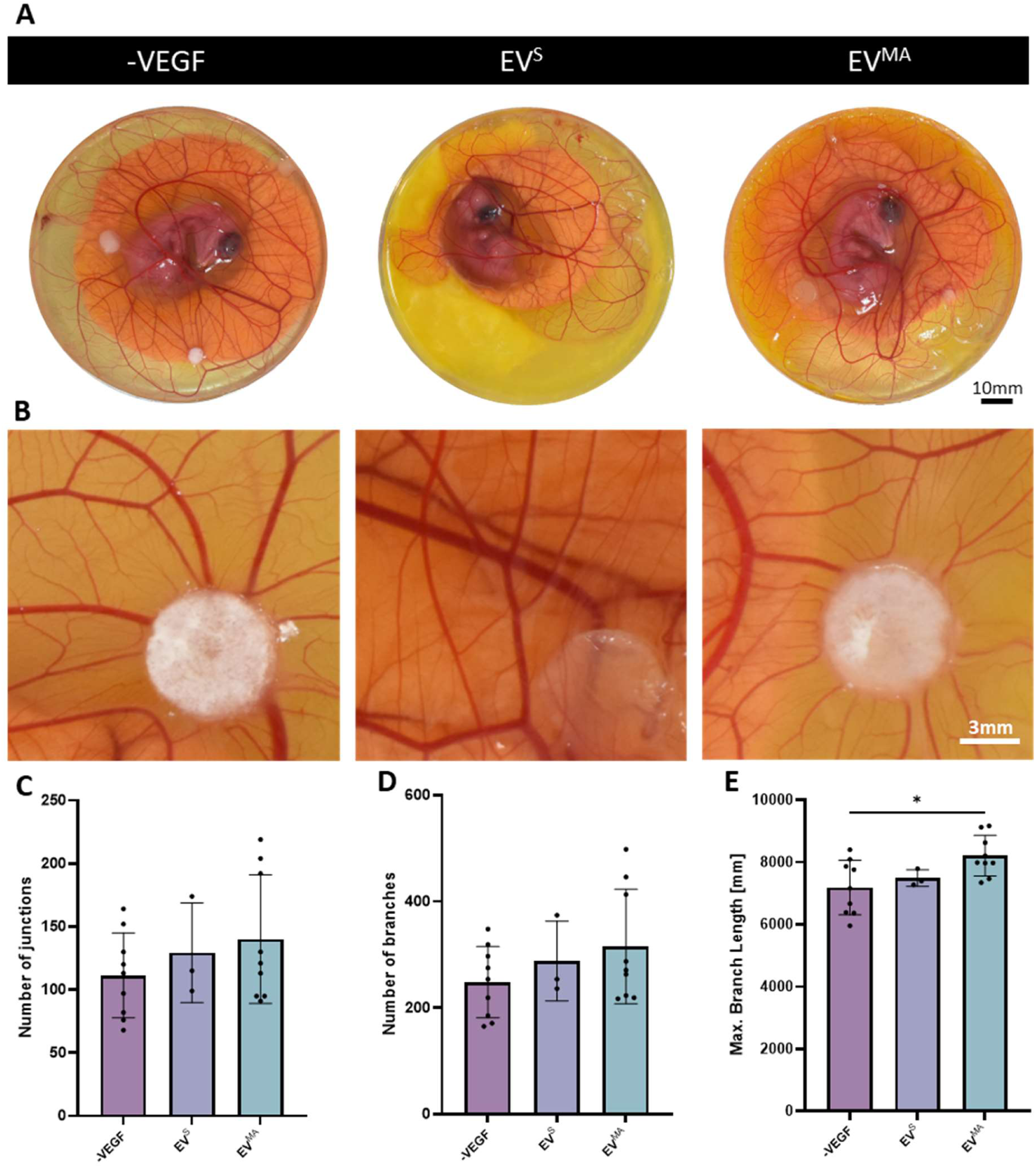
Effect of static and mechanically activated OCY-EVs on angiogenesis in a chick embryo CAM *ex ovo* model. **(A)** Representative images of the angiogenic response of the EVs on the CAM **(B)** Selected zones for vessel formation analysis **(C)** Number of junctions **(D)** Number of branches **(E)** Maximum branching length. Significant differences reported when tested using a one-way ANOVA. Data presented as Mean ± SD, n≥3.

## Discussion

The formation of new blood vessels, angiogenesis, is vital for both bone homeostasis and repair. This process is highly dependent on support cell populations and by the biochemical and biophysical cues of the cellular environment. The overall goal of this study was to assess the effect of the bone cell secretome and associated EVs as they differentiated through the osteogenic lineage on angiogenesis, as well as the impact of mechanical environment on this response. Therefore, we have established for the first time a relationship between the stage of osteogenic lineage and the angiogenic impact of mechanically-stimulated bone cell-derived EVs. MSCs, OBs and OCYs were placed in a PPFC bioreactor and both their static and mechanically-activated secretomes and associated EVs were collected and used to treat HUVECs, showing differences according to the stage of lineage and particularly the application of mechanical stimulation of the parent cell. In particular, OCY-derived EVs demonstrated superior efficacy and were further shown to enhance endothelial cell migration, increase CD31 expression, and promote the formation of more complex tube structures employing the optimal dose of 0.5 µg/mL, as shown by a dose response study. Lastly these EVs demonstrated superior vessel formation compared to the PBS control, in a preclinical *ex ovo* CAM model. Therefore, the mechanically activated OCY-derived EVs represent an innovative and powerful pro-angiogenic tool to promote vessel formation.

The secretome of MSCs has been substantially investigated for its regenerative properties, as it has been shown to modulate angiogenesis, immune response, and extracellular matrix remodelling [28, 42–44]. Focusing on angiogenesis, a plethora of previous studies have shown favourable results with MSC-derived secretome [28, 45–47]. For example, Estrada *et al*. found that MSC-derived secretome increases HUVEC tube formation; though, high doses (minimum 5 µg/mL) of filtered murine derived secretomes were used [28]. On the contrary, we have shown that hMSCs do not secrete factors that influence angiogenesis, at least under the conditions of our study. Remarkably, when these same cells undergo mechanical stimulation, they secrete paracrine factors that promote angiogenesis. These paracrine factors within the secretome may be packaged in EVs or circulate unbound in the media. Different growth factors and cytokines are known to be present, such as VEGF and TGF-*β*1 that have been shown to be released by MSCs and promote angiogenesis by activating the PI3K/Akt and MAPK signalling pathways [47–49]. Interestingly, the EVs obtained from both the CM^S^ and CM^MA^ had no effect on angiogenesis, suggesting that the effects seen with CM-treatments are likely due to soluble factors in the media. Several studies have investigated the role of MSC-EVs on angiogenesis[50–55], where increases in migration, proliferation and tube formation were seen with EV treatments, although with significantly higher EV doses (i.e. 10 µg/mL, 160 µg/mL) which can be challenging to scale-up commercially.

We then developed a protocol to differentiate MSCs into OBs and lift these cells from their matrix, so that they could be reseeded in our bioreactor in a similar manner to that achieved with MSCs. The coupling of osteoblasts and endothelial cells within the angiogenic-osteogenic cell niche makes the secretome and EVs of these cells appealing for angiogenic bone repair focused therapies. Particularly, osteoblasts and endothelial cells communicate through Notch signalling to regulate bone homeostasis [56]. CM^S^ from these cells had no effect on angiogenesis, similarly to the MSC-CM^S^. Interestingly, the same donor of MSC differentiated into OBs showed different angiogenic potency results, with a loss of effect with the mechanically stimulated group. Nevertheless, the effect of a mouse osteoblastic cell line derived secretome has been assessed and shown similar results to our study [34]. Specifically, no differences were seen in tube formation between static and mechanically activated secretomes in both studies. Focusing on OB-EVs derived from this secretome, we saw an increase in tube formation with EV^MA^ group, which may be related to the mechanical environment that osteoblasts are experiencing in the angiogenic-osteogenic niche, allowing them to secrete EVs with pro-angiogenic cargos. This is the first time the effect of human OB-EVs on angiogenesis has been studied, as well as the effect of mechanical stimulation on the angiogenic effect of these derived EVs. A previous study from our group reported similar results with mechanically-activated EVs derived from the murine osteoblastic cell line MC3T3-E1, which further confirms the possible use of osteoblast derived EV^MA^s as a treatment for bone repair [26].

EV^MA^s collected from the final stage of the mesenchymal lineage, OCYs, had an optimal effect on angiogenesis, with significant increases in all tube formation readouts, a response that was not seen with EV^S^. To ensure the optimal EV dose, a dose response study was performed, where the same concentration of EV^MA^s that was present in the CM (0.5 µg/mL) had similar number of junctions and higher number of segments when compared to higher doses of 2 µg/mL and 4 µg/mL. This suggests that the therapeutic potential of EVs is dose dependent. Hagey *et al.* confirmed this hypothesis by verifying that the cellular response to EVs is not only cell source dependent but also dose dependent [57]. Moreover, they found that the greatest responses were observed when EVs were applied at low doses, similarly to the results seen in this study. Since the 0.5 µg/ml dose exhibited enhanced angiogenic potency as well as lower variability compared to the other concentrations, it was chosen as the optimal dose for subsequent studies. Considering that angiogenesis begins with migration and proliferation, followed by the formation of tubular structures [58, 59], both processes were assessed to have a complete view of the angiogenic potency of these EVs. While, OCY-EVs had no effect on the proliferation of HUVECs, EV^MA^ treatment led to an enhancement in migration. Platelet endothelial cell adhesion molecule (PECAM-1) or CD31 is a vascular cell adhesion and signalling molecule that is highly expressed in endothelial cells [60]. Treatment with OCY-EV^MA^s showed increased CD31 expression, indicating more mature tubule formation [61]. This result could indicate that these EVs lead to vessel formation more in line with type H vessels (highCD31/ high EMCN), which are vital in normal bone functioning and regeneration [62]. Moreover, CD31 also has a role in endothelial cell migration, which correlates to the results seen with the migration assay [60]. We have shown for the first time a link between mechanically activated osteocyte secretome and the expression of CD31 in endothelial cells. Having demonstrated the potential of OCY-EVs *in vitro*, we next assessed these EVs in a more physiologically relevant model, by utilising a chicken embryo *ex ovo* CAM assay, which provides a vascularised microenvironment to study angiogenic and osteogenic responses *in vivo*. To the best of our knowledge, this is the first report of OCY-EVs analysed in an angiogenesis chick embryo model. EV^MA^s treatment led to an increase in total tube length of the microvasculature in the CAM, which was congruent with the results seen *in vitro*.

A possible limitation of this study is the use of both human (MSC and OB) and murine (OCY) derived EVs to treat human cells. However, despite this shift in species, the trends of behaviour are consistent indicating that the MLO-Y4 may represent a suitable model for human osteocytes[63–65]. Moreover, from a clinical perspective, the use of non-human derived EVs may raise concerns over potential adverse immune responses. However, several studies have assessed the effect of EVs used to treat animals of different species with no adverse effects reported [66–69]. Furthermore, EVs possess several mechanisms that allow them to avoid immune interactions, such as the surface protein CD47, which allows them to avoid the mononuclear phagocytic system (MPS)[70]. In conclusion, we have for the first time elucidated the role of mechanical stimulation and parent cell lineage stage on the angiogenic properties of bone cell derived secretomes and associated EVs. Furthermore, we have highlighted the potential therapeutic benefit of mechanically-activated OCY-EVs as a novel angiogenic therapy tailored for bone repair.

## Acknowledgments

Funding was provided by Research Ireland through the Frontiers for the Future Project [19/FFP/6533] and Award [23/FFP-A/12166], by EU Horizon 2020 IND, EVPRO to LOD [814495], by the Irish Research Council Advanced Laureate Award EVIC to LOD [IRCLA/2019/49], by the Research Ireland Enterprise Partnership Scheme award [EPSPD/2024/676].

